# Counterconditioning Alcohol Cues: Neural and Behavioral Modulation of Automatic Tendencies and Pavlovian-to-Instrumental Transfer in Male Alcohol Users

**DOI:** 10.1101/2025.11.10.687519

**Authors:** Adarsh K. Verma, Usha Chivukula, Maria Garbusow, Neeraj Kumar

**Affiliations:** Department of Psychology, University of Hyderabad, India; Department of Psychology, Medical School Berlin, Germany; Department of Liberal Arts, Indian Institute of Technology Hyderabad, India

**Keywords:** Alcohol use behavior, automatic tendencies, Pavlovian-to-Instrumental Transfer (PIT), event-related potentials, counterconditioning

## Abstract

**Objective:** Repeated alcohol-cue pairings with rewarding experiences establish automatic response tendencies that bias instrumental behavior through Pavlovian-to-instrumental transfer (PIT). How individual differences in approach and avoidance tendencies shape PIT and intervention responsiveness remains poorly understood. This study examined whether automatic tendencies differentially modulate PIT, and whether counterconditioning (CC) combined with retrieval cues can attenuate approach biases and cue-driven instrumental interference.

**Method:** Thirty-nine male alcohol users completed an Alcohol Approach-Avoidance Task and PIT task before and after CC, classified into approach or avoidance groups based on their baseline Alcohol Approach Index. Alcohol cues were paired with monetary loss during CC, with retrieval cues introduced to facilitate reactivation of newly formed aversive associations. Right frontal N2 and centroparietal P3 were examined as indices of early evaluative and attentional processing.

**Results:** Despite comparable behavioral PIT performance, approach-oriented individuals showed attenuated right frontal N2 during push actions at baseline, reflecting reduced frontal regulatory engagement during cue-incongruent responding. CC selectively reduced approach biases in the approach group, with retrieval cues producing additional reductions, accompanied by N2 and P3 restoration. CC recalibrated PIT interference by reducing push and increasing pull interference, particularly with retrieval cues. CC-induced changes in approach bias did not predict PIT interference, suggesting modulation through partially dissociable pathways.

**Conclusions:** CC with retrieval cues selectively reduced approach biases and recalibrated cue-driven instrumental interference in approach-oriented individuals, providing proof-of-concept for retrieval cue-integrated CC as a targeted associative intervention and highlighting the importance of individual tendency profiles in intervention responsiveness.

## Introduction

Repeated pairings of alcohol cues with positive experiences establish robust associative memories that, over time, manifest as automatic approach tendencies operating below conscious awareness (Drummond et al., 1990; Lutz & Childs, 2021). When encountered in everyday contexts, such as passing a wine shop while commuting, these cues can interfere with intended behavior and undermine adaptive control, even in individuals motivated to abstain (Abiero et al., 2022; Campanella et al., 2019; Field et al., 2008; Garbusow et al., 2022; Wiers & van Dessel, 2023). One mechanism through which this occurs is Pavlovian-to-instrumental transfer (PIT), whereby Pavlovian cues (e.g., appetitive alcohol cues) modulate instrumental action selection toward cue-congruent responses without directly signaling which action is most effective (Garbusow et al., 2019; Hogarth et al., 2014, 2019). Debate continues as to whether PIT reflects habitual or goal-directed control, with some evidence suggesting cue-elicited responding operates independently of outcome value (Hogarth & Chase, 2011; Watson et al., 2014), while other studies demonstrate sensitivity to outcome devaluation supporting goal-directed mechanisms (Hogarth et al., 2019; Seabrooke et al., 2019). Importantly, these investigations have predominantly focused on group-level effects, treating individual differences in automatic tendencies as noise rather than as a theoretically meaningful source of variance.

Considerable heterogeneity exists in how individuals respond to alcohol cues, with some showing strong approach tendencies while others exhibit avoidance patterns despite continued drinking (Kenney et al., 2018; McNaughton et al., 2016; Morris et al., 2020; Verma et al., 2025). Whether these opposing automatic tendencies produce distinct patterns of cue-driven interference with instrumental responding remains largely unexplored. If approach and avoidance tendencies differentially modulate PIT, this would suggest that cue-driven interference is not uniform but shaped by pre-existing motivational biases, with implications both for theoretical accounts of habitual versus goal-directed control and for the design of targeted interventions.

Interventions developed to reduce automatic approach biases, including cognitive bias modification, avoidance training, and behavioral extinction, show important limitations, with effects that are often short-lived and poorly generalized beyond controlled settings (Bouton, 2002; Cristea et al., 2016; Rinck et al., 2018; Van Gucht, Van den Bergh, et al., 2010; Wiers et al., 2011, 2013). A major reason behind this limited success is context dependency, whereby benefits observed during training do not transfer to real-world drinking situations. This underscores the need for interventions that target both the reduction of maladaptive automatic tendencies and the restoration of goal-directed control, while accounting for contextual generalization. Counterconditioning (CC) represents a theoretically grounded alternative. Rather than suppressing approach tendencies, CC creates competing associative memories by pairing alcohol cues with aversive outcomes, thereby reassigning their motivational value and weakening their capacity to bias instrumental behavior (Baker & Cannon, 1979; Das et al., 2015; Goltseker et al., 2016, 2021). This is conceptually consistent with Hogarth et al.’s (2014) demonstration that degrading cue-outcome expectancies, rather than simple extinction, effectively abolished PIT effects. CC may operate through a similar mechanism by altering the hierarchical signaling function of alcohol cues. Critically, the effectiveness and generalizability of CC may be further enhanced by incorporating retrieval cues, neutral salient stimuli presented during CC training and reintroduced at test to reactivate the newly formed aversive associations across contexts, supporting generalization of CC effects beyond the training environment (Collins & Brandon, 2002; Stasiewicz et al., 2007; Treanor et al., 2017).

The present study first examined whether automatic approach and avoidance tendencies differentially modulate PIT, and subsequently assessed whether CC combined with retrieval cues reduces approach biases and alcohol-related PIT interference at both behavioral and neural levels. To capture the interplay between automatic and controlled processes underlying these effects, we assessed neural activity at frontal and centroparietal regions implicated in cue-driven cognitive regulation (da Silva et al., 2013; McNeill et al., 2022). Frontal N2 reflects conflict-related processing arising from competing cue-driven and instrumental action tendencies (Buss et al., 2011; Espinet et al., 2012), while centroparietal P3 indexes controlled attentional processing of motivationally relevant stimuli (Polich, 2007; Reed et al., 2022; Sawaki & Katayama, 2008). We expected individuals with approach tendencies to show greater cue-driven interference during instrumental responding, particularly for avoidance actions in the presence of appetitive alcohol cues. We further hypothesized that CC would reduce automatic approach biases toward alcohol cues as measured by the AAT, disrupt the learned associations sustaining cue-driven PIT interference, and enhance neural markers of cognitive control, with retrieval cues facilitating the generalization of these effects beyond the training context. The current paradigm tests fundamental principles of how CC modulates alcohol approach tendencies and their influence on instrumental behavior, with findings intended to inform the development of future multi-session clinical interventions.

## Method

### Sample

A priori power analysis was conducted using G*Power 3.1 (Faul et al., 2007) to estimate the required sample size for a repeated measures design with a within–between interaction. The analysis was based on an assumed moderate effect size (f = 0.25), an alpha level of 0.05, and a statistical power of 0.80. The analysis indicated that in total, a minimum sample size of 24 participants would be required to detect the expected effects.

To ensure sufficient representation of both approach and avoidance subgroups, 40 alcohol users were recruited. All participants were the same gender (males), which facilitated reducing the variance associated with gender differences in alcohol use patterns and approach–avoidance responding, thereby increasing internal validity for mechanistic investigation.

During statistical analysis, one participant’s data was dropped due to large EEG artifacts. The remaining 39 participants’ (mean age = 21.7 years, SD = 2.32) data were included in the analysis. All participants reported being involved in alcohol consumption and were screened using the Alcohol Use Disorders Identification Test (AUDIT; Saunders et al., 1993), where 27 participants’ scores ranged from 1 to 7 (indicating low-risk alcohol consumption) and 12 participants’ scores ranged between 8 to 14 (indicating harmful alcohol consumption habits). At the time of participation, none of the participants met the diagnostic criteria for Alcohol Use Disorder (AUD). All participants were right-handed or mixed right-handed, as assessed using the Edinburgh Handedness Inventory-SF (Veale, 2014), and had no history of neurological conditions. Post-A-AAT administration, 28 participants were classified into the alcohol approach group (having a positive approach-avoidance index [AAI] on a continuous scale) and 11 into the alcohol avoidance group (negative AAI) for statistical analysis (detailed classification reported in Data Analysis section).

All participants were informed in advance to abstain from alcohol consumption for a minimum of 24 hours prior to the experimental session. The study adhered to the ethical guidelines of the Declaration of Helsinki and was approved by the relevant Institutional Ethics Committees [*details blinded for review*]. Written informed consent was obtained from all participants prior to the commencement of the study.

### Experimental Design and Tasks

The experiment was programmed in MATLAB using the Psychophysics Toolbox version 3.0.19 (Brainard, 1997) and presented on a 27-inch, 120 Hz monitor in portrait orientation. All tasks were completed in a dark, sound-attenuated room to minimize distractions and maintain consistency. The study followed an ABA design: participants first completed the Alcohol Approach-Avoidance Task (A-AAT) and Pavlovian-to-Instrumental Transfer (PIT) task in Context A (teal background), followed by a counterconditioning (CC) phase in Context B (gray background), and then repeated the A-AAT and PIT tasks in Context A (Figure 1A) (Stasiewicz et al., 2007; Torregrossa & Taylor, 2012). CC was implemented in this distinct context to test generalization beyond the training context, with retrieval cues integrated within the counter-conditioning protocol to enhance the transfer of interventional effects across contexts. These context shifts simulated real-world treatment generalization, mimicking transitions between clinical and everyday settings. A five-minute break was provided between tasks to reduce carryover effects.

**Figure 1.**
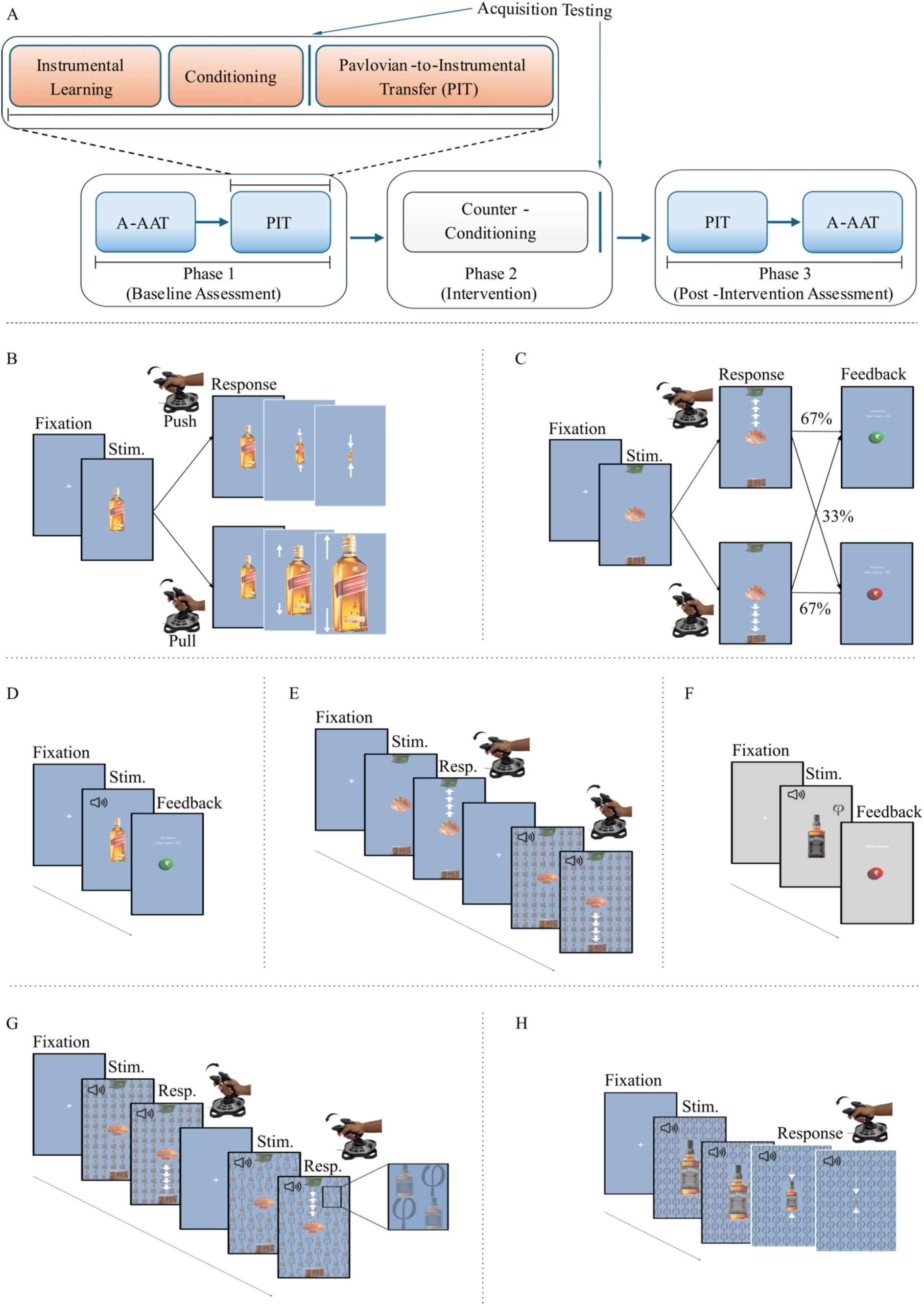
Overview of the experimental paradigm. The study followed an ABA design, with tasks administered across three phases: baseline, pre-counterconditioning (pre-CC), and post-counterconditioning (post-CC) (Panel A). In the Approach–Avoidance Task (A-AAT, Panels B and H), participants used a joystick to pull (approach) or push (avoid) alcohol and non-alcohol stimuli. Movements triggered a simulated zoom-in or zoom-out effect along the y-axis, though these visual effects were not visible to participants. Panel C depicts the instrumental training phase, during which participants learned through trial and error to pull “good” shells toward a chest and push “bad” shells toward a bin. Panel D illustrates Pavlovian conditioning, in which alcohol-related images were repeatedly paired with monetary rewards. Panel E displays the pre-CC Pavlovian-to-Instrumental Transfer (PIT) task, in which background alcohol cues modulate instrumental responses. During counterconditioning (Panel F), alcohol cues were paired with negative outcomes, including monetary loss, with and without symbolic retrieval cues. Panel G presents the post-CC PIT task, where symbolic retrieval cues were reintroduced to assess their effect on transfer learning. Finally, Panel H shows the post-CC A-AAT, used to evaluate changes in automatic action tendencies. The white arrows shown in the figure (in Panels B, C, E, G, and H) were not visible to participants and are included solely to illustrate movement direction and cue dynamics. NR = no retrieval-cue trials post-CC, R = trials with a retrieval cue post-CC

#### Alcohol Approach-Avoidance Task (A-AAT)

The adapted A-AAT was used to measure automatic approach or avoidance towards alcohol-related stimuli (Figure 1B) (Korucuoglu et al., 2016; Rinck & Becker, 2007). It included 180 trials divided into three blocks: two experimental blocks (80 trials each) and a neutral washout block (20 trials). In Block 1, participants were instructed to pull the joystick in response to alcohol images (approach) and push it for non-alcohol images (avoidance). Pulling or pushing resulted in a dynamic zoom-in or zoom-out effect, respectively, proportional to the joystick displacement along the y-axis. Alcohol and non-alcohol images were presented on 50% of trials each, with the sequence programmed to prevent more than three consecutive presentations of the same image type. In the washout block, eight neutral images (e.g., bells and light bulbs) were used, where participants performed similar joystick movements (pulling bells and pushing light bulbs) to minimize any motor learning biases from Block 1. In Block 2, the instructions were switched, i.e., participants now pushed alcohol images and pulled non-alcohol images.

Each trial began with a 500-ms fixation cross, during which participants were required to maintain a neutral joystick position for at least 200 ms to prevent unintentional/premature responses. The stimulus then appeared for a maximum of 3000 ms, during which participants executed the required response. During practice trials, incorrect responses were flagged with a red cross, while a green tick indicated correctness. During the main trials, only feedback for incorrect responses was shown to help correct future trials.

#### Pavlovian-to-Instrumental Transfer (PIT) Task

The PIT paradigm consisted of three phases: instrumental training, Pavlovian conditioning, and transfer testing (adapted from Belanger et al., 2022; Garbusow et al., 2019). It was employed to assess whether Pavlovian cues modulate ongoing instrumental behavior.

##### Instrumental Learning

In an instrumental learning phase, participants completed 96 trials to learn, through trial-and-error, the reward or loss probabilities associated with six different shell stimuli (varied in color, shape, and texture). Each trial starts with a 500 ms fixation, followed by a 2000 ms stimulus screen displaying a colored shell at the center, a treasure chest at the bottom, and a trash can at the top (Figure 1C). Randomized across subjects, three shells were designated as “*good*” and assigned a positive value, while the remaining three were “*bad*” and assigned a negative value. Participants collected shells by pulling the joystick toward the treasure chest and rejected shells by pushing the joystick toward the trash can. Collecting a *good* shell produced a reinforcement of 50 tokens (gain), whereas rejecting it resulted in a punishment of 50 tokens (loss), with the reverse contingencies for *bad* shells. Each shell type appeared in 50% of trials, and approximately one-third of trials pseudo-randomly reversed the expected outcome to maintain uncertainty and prevent perfect memorization. Feedback indicating a gain or loss of 50 tokens was presented for 1500 ms after each response. Through this phase, participants acquired instrumental responses by learning the action–outcome contingencies associated with different shell types, enabling them to select the optimal action (collect vs. reject) to maximize token gains. For each trial, response accuracy (correct vs. incorrect), joystick position, and timestamps were recorded. Movement velocity was computed offline as the rate of change in joystick position over time. Participants were informed that the monetary compensation at the end of the experiment would be proportional to the earned tokens.

##### Pavlovian Conditioning

For the Pavlovian conditioning phase, participants completed 100 trials to establish stimulus–outcome associations. Each trial began with a 3000-ms compound audiovisual cue, consisting of either an alcohol or a non-alcohol image presented at the center of the screen along with a specific tone delivered through headphones. This was followed by a 3000-ms outcome screen (Figure 1D). In 80% of trials, alcohol cues were paired with positive outcomes (e.g., +100 tokens) and non-alcohol cues with negative outcomes (e.g., –100 tokens), while in the remaining 20% of trials these contingencies were reversed. Following conditioning, an acquisition test assessed participants’ explicit knowledge of the learned cue–outcome contingencies by requiring them to indicate whether each image–tone pair had been associated with reward or punishment. The accuracy of their responses was recorded.

##### PIT Phase (Pre-CC)

This phase evaluated whether Pavlovian cues influenced instrumental decisions, with the task structure similar to the instrumental phase (Figure 1E). The task consists of 120 trials. Each trial began with a 500 ms fixation, followed by a 2000 ms stimulus screen. In 80% of the trials, the shell stimuli were presented against a tiled background containing the previously conditioned alcohol cues and associated tone, and the remaining 20% against non-alcohol cues and respective tone. In alcohol congruent trials (40%), positively conditioned alcohol cues appeared with *good* shells; in alcohol incongruent trials (40%), they appeared with *bad* shells. During the task, no feedback was provided to prevent the formation of new cue-outcome contingencies, but participants were told their decisions still affect their total token earnings.

#### Counterconditioning (CC)

The CC phase was conducted to reverse previously formed positive associations between alcohol cues and reward (Furlong et al., 2015; Stasiewicz et al., 2007). Across 120 trials, alcohol images were paired with monetary losses under varying stimulus conditions (Figure 1F). Specifically, in 80% of the trials, alcohol images with the respective tone were followed by loss outcomes, either a loss of 100 tokens or a larger loss of 500 tokens when accompanied by a retrieval cue (a Greek symbol, φ). The larger loss magnitude when the retrieval cue was present was designed to enhance the salience and memorability of the retrieval cue-alcohol association, though we acknowledge this may have also allowed the retrieval cue to form direct Pavlovian associations with loss (see Limitations).

To maintain learning variability and introduce decision uncertainty, 20% of the trials retained prior reward contingencies. These included audiovisual alcohol stimulus resulting in a gain of 100 tokens, or with another cue (ϰ), leading to a larger gain of 500 tokens. The CC task was administered in Context B (gray background) to simulate a different setting in which interventions are typically delivered, thereby promoting contextual separation and supporting the generalization of new learning. Retrieval cues, visible only on screen and not explained to participants, were intended to later serve as symbolic reminders of the counterconditioning context during subsequent testing. Each trial began with a 500 ms fixation, followed by a 3000 ms stimulus display and a 3000 ms outcome screen. Similar to the conditioning phase, participants completed an acquisition test after CC to evaluate the newly formed cue–outcome associations.

#### Post-CC PIT and A-AAT

In the Post-CC PIT, participants completed a structurally identical task to the pre-CC PIT (Figure 1G). In this phase, the background grids contained counterconditioned alcohol images in 80% of trials. In congruent trials, negatively conditioned alcohol cues accompanied *bad* shells, while in incongruent trials, they were presented with *good* shells. Among the trials containing counterconditioned stimuli, half included retrieval cues presented in an alternate grid (Figure 1G), while the other half were presented without retrieval cues, allowing for the examination of the influence of symbolic reminders on behavior.

In the Post-CC A-AAT, the task structure remained identical to the original A-AAT, but counterconditioned alcohol stimuli were used (Figure 1H). Each image was randomly presented with or without a retrieval cue, in 50% of trials each. This manipulation allowed assessment of the impact of the retrieval cue on approach and avoidance responses.

#### Compensation and Debriefing

At the completion of the experiment, participants were monetarily compensated based on their performance, with a fixed minimum amount, and fully debriefed about the experiment.

### EEG Data Acquisition and Processing

EEG data were recorded using the Brain Products ActiChamp Plus system with 64 Ag/AgCl electrodes at a sampling rate of 1 kHz. During preprocessing, a 50 Hz notch filter was applied to remove powerline interference. A bandpass filter (0.1–30 Hz) was then applied, followed by epoching (from −1000 to 2000 ms), rejection of artifactual trials, and independent component analysis (using the runica algorithm) to remove eye blink and muscle artifacts. A baseline correction period of −200 to 0 ms relative to stimulus onset was applied to all epochs. All preprocessing was conducted using the EEGLAB toolbox in MATLAB (Delorme & Makeig, 2004).

The ERPs of interest, i.e., N2 at frontal regions and P3 at parietal regions, were extracted based on prior literature. To optimize signal clarity of concerned ERPs, a 0.2–10 Hz bandpass filter was applied (Zhang et al., 2024). The ERP analyses targeted electrode clusters over the left and right frontal regions and the centro-parietal region. The left frontal region of interest (LF) was defined as the average activity from electrodes F1, F3, F5, F7, AF3, FC3, and FC5. The right frontal region of interest (RF) was defined using electrodes F2, F4, F6, F8, AF4, FC4, and FC6. The centro-parietal region of interest (CP) was the average of CPz and Pz (Sochurková et al., 2006; Zhang et al., 2024). The mean amplitude of N2 was computed as the average amplitude between 250–650 ms post-stimulus at LF and RF, while P3 was extracted from CP in the 350– 650 ms window based on a visual inspection of grand-averaged waveforms.

### Data Analysis

Response time (RT) in the A-AAT was defined as the time elapsed between stimulus onset and maximum joystick displacement (±1). For each participant, automatic approach scores were calculated by subtracting median pull RT from push RT separately for alcohol and non-alcohol images. The Alcohol Approach Index (AAI) was then derived by subtracting the non-alcohol score from the alcohol score, with positive values indicating approach bias and negative values indicating avoidance bias. Group comparability on demographic characteristics was assessed using Wilcoxon rank-sum or Chi-square tests, and instrumental learning was operationalized as ≥80% accuracy in the last 16 trials (Belanger et al., 2022). Behavioral and neural data were analyzed using parametric ANOVAs, with ART-ANOVAs applied when assumptions were violated (Wobbrock et al., 2011), followed by post-hoc Wilcoxon or t-tests. CC effects were assessed by comparing baseline, post-CC without retrieval cue, and post-CC with retrieval cue conditions across A-AAT, PIT, and ERP outcomes (N2 at RF; P3 at CP).

Spearman correlations examined associations between AAI and ERP alcohol–non-alcohol difference scores, and between action-related ERP differences and PIT interference scores, with changes in correlation strength assessed using Steiger’s tests (Steiger, 1980). To examine whether CC-induced PIT changes were independently predicted by concurrent changes in approach tendencies, PIT change scores were regressed on AAI change scores separately for each retrieval cue condition, with robust HC3 standard errors applied as a robustness check. All analyses were conducted in R using ARTool, wilcox_effsize, and psych packages, with Benjamini–Hochberg corrections for multiple comparisons and Jeffreys–Zellner–Siow Bayes factors (r = 0.707) computed for all null results.

Although LF ERP amplitudes were analyzed, all comparisons at this site were non-significant across tasks and are not reported. The non-significant results are consistent with LF’s established role in craving-related processing rather than rapid cue-action regulation, as shown in previous substance use research (da Silva et al., 2013; McNeill et al., 2022).

## Results

The current study examined how automatic tendencies towards alcohol influence instrumental behavior in the presence of appetitive alcohol cues. Additionally, the study explores how counterconditioning (CC) can modulate the automatic alcohol approach tendencies and PIT.

### Preliminary Analysis

Preliminary analyses show similar demographic characteristics in the Approach and Avoidance groups, i.e., no significant differences in age (*W* = 93, *p* = 0.056), AUDIT scores (*W* = 115.5, *p* = 0.233), or education level (Undergraduate / Postgraduate / Doctorate: *χ²*_(2, N = 39)_ = 1.31, *p* = 0.520). Additionally, the distribution of participants across AUDIT risk categories (low, moderate, and high) did not differ significantly between groups (*χ²*_(2, N = 39)_ = 4.38, *p* = 0.112). Descriptive statistics for all variables are presented in Table 1.

**Table 1.** Demographic and baseline characteristics of Approach and Avoidance groups. AUDIT scores are categorized according to drinking patterns: low-risk (1–7), moderate-risk (8–14), and high-risk (15+).

|  |  | Approach | Avoidance |
| --- | --- | --- | --- |
| N [% of total] |  | 28 [71.79] | 11 [28.21] |
| Age (mean ± SD) | | 21.21 ± 2.01 | 22.82 ± 2.75 |
| AUDIT (n [% within group]) | low risk (1-7) | 21 [75.00] | 5 [45.45] |
|  | moderate risk (8-14) | 4 [14.29] | 5 [45.45] |
|  | high risk (15+) | 3 [10.71] | 1 [9.09] |
| AAI (Mean ± SE) | | 0.125 ± 0.018 | -0.081 ± 0.025 |
| Educational program (n [% within group]) | Undergraduate | 17 [60.71] | 5 [45.45] |
|  | Postgraduate | 9 [32.14] | 4 [36.36] |
|  | Doctorate | 2 [7.14] | 2 [18.18] |

### Baseline Assessment: Approach Tendencies Shape Neural Responses to Alcohol Cues

#### Approach Tendencies Elicit Attenuated Frontal–Parietal Neural Response to Alcohol Cues During A-AAT

During baseline assessment, participants were divided into approach and avoidance groups based on their Alcohol Approach Index (AAI) (Figure 2A). To rule out a generic motor bias (i.e., faster pull than push responses regardless of stimulus type), response times were analyzed across Stimulus (alcohol vs. non-alcohol), Movement (pull vs. push), and Group (approach vs. avoidance). A significant three-way interaction emerged (*F*_(1,111)_ = 65.79, *p* < 0.001), indicating that modulation of pull and push responses differed by group and alcohol. In the approach group, pull responses were faster for alcohol than non-alcohol stimuli (median difference = −0.069, *W* = 0, *p* < 0.001, *r* = 0.874), but slower for push responses (median difference = 0.043, *W* = 372, *p* < 0.001, *r* = 0.727). In the avoidance group, alcohol–non-alcohol differences were absent for pull responses (median difference = 0.025, *W* = 53, *p* = 0.083, *BF_10_* ≈ 0.219) but significant for push responses (median difference = −0.025, *W* = 3, *p* = 0.014, *r* = 0.765). These results confirm that AAI-based classification captures stimulus- and action-specific processing of alcohol cues, not a generalized pull–push motor bias.

**Figure 2.**
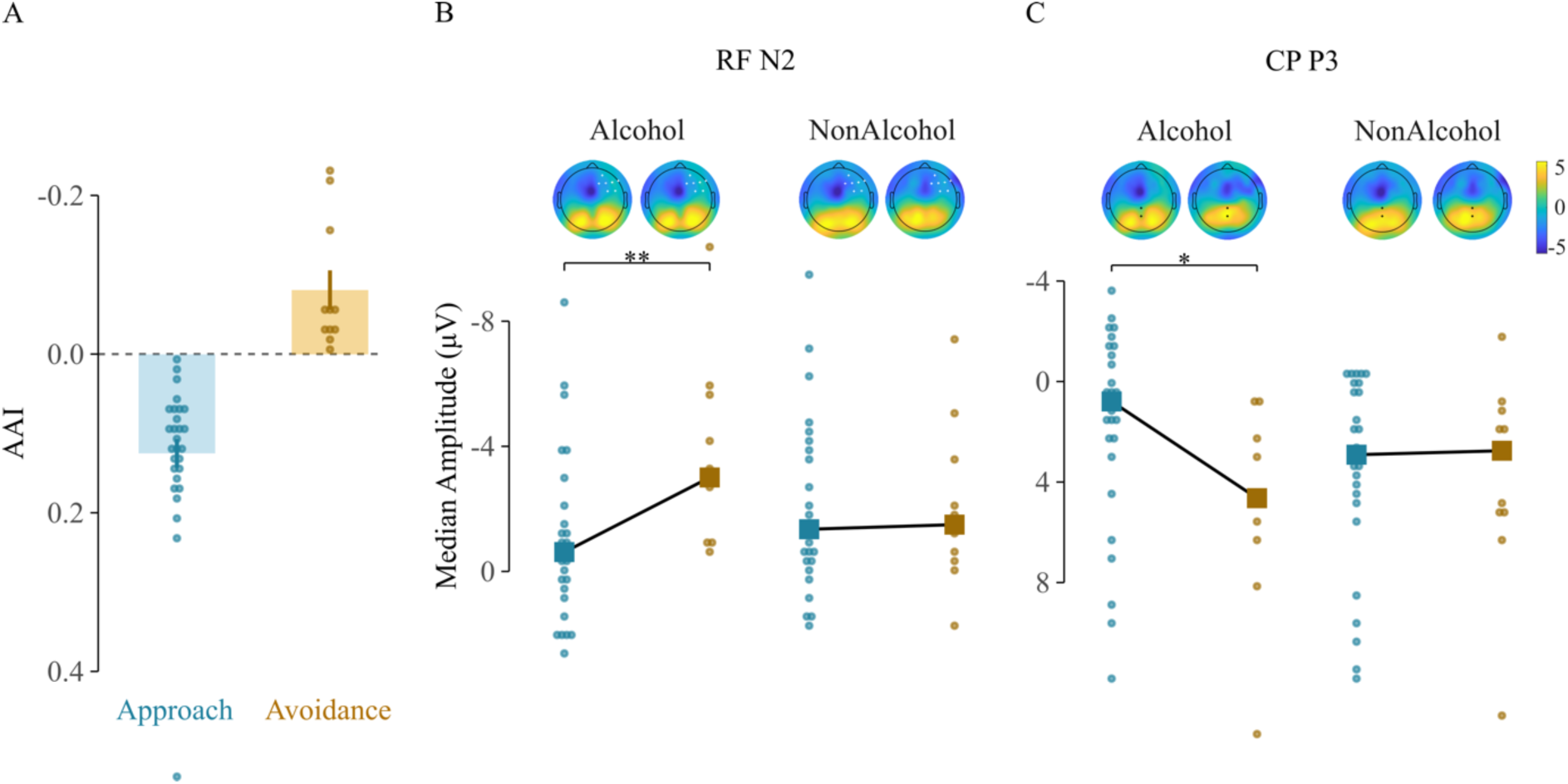
Automatic alcohol approach tendencies compromise neural responses to alcohol stimuli. Panel A depicts individuals classified into approach and avoidance groups based on the Alcohol Approach Index (AAI). At both RF (Panel B) and CP (Panel C), individuals with approach tendencies show significantly attenuated neural responses to alcohol cues compared to those with avoidance tendencies. Blue and brown colors indicate approach and avoidance group data points, respectively. AAI = Alcohol Approach Index; RF = right frontal region; CP: centro-parietal region. * p < 0.05; ** p < 0.01

The validity of the AAI-based classification was further supported by the ERP data. Approach and avoidance groups differed in neural responses to alcohol cues, independent of movement direction (Stimulus × Movement × Group interaction: RF, *F*_(1,111)_ = 0.034, *p* = 0.855, *BF_10_* ≈ 0.032; CP, *F*_(1,111)_ = 0.983, *p* = 0.324, *BF_10_* ≈ 0.017). A significant Stimulus × Group interaction was observed at right-frontal sites (*F*_(1,111)_ = 51.212, *p* < 0.001; Figure 2B) and centroparietal sites (*F*_(1,111)_ = 39.493, *p* < 0.001; Figure 2C). The approach group exhibited attenuated N2 (median difference = 2.39, *W* = 244, *p* = 0.004, *r* = 0.461) and P3 (median difference = –3.850, *W* = 70, *p* = 0.008, *r* = –0.425) amplitudes to alcohol relative to non-alcohol stimuli, consistent with reduced engagement of cognitive control mechanisms. Within the approach group, alcohol-specific attenuation was evident for RF N2 (median difference = 0.148, *W* = 500, *p* = 0.078, *r* = 0.236) and CP P3 (median difference = –2.120, *W* = 264, *p* = 0.036, *r* = –0.280). Together, these findings indicate that the AAI-based classification reflects stimulus-specific neural processing differences rather than a generalized pull–push motor bias.

#### Automatic Approach Tendencies Attenuate Neural Control During Cue-Incongruent Push Actions During PIT

To examine how automatic approach and avoidance tendencies influence instrumental behavior, all participants completed the PIT task. During the instrumental learning phase, both groups successfully learned the instrumental contingencies linking specific shell stimuli with reward or loss outcomes (approach participants: 81.2% accuracy; avoidance participants: 86.4% accuracy), as reflected in the last 16 trials (Belanger et al., 2022).

To assess alcohol cue interference during the PIT phase, error rates were examined as a function of Group (Approach vs. Avoidance) and Action (Pull vs Push). Overall error rates were comparable across approach and avoidance participants (*F*_(1,74)_ = 0.906, *p* = 0.344), indicating that alcohol cues elicited similar levels of Pavlovian interference in both groups. However, performance differed across participants depending on whether the required instrumental response was pull (alcohol-congruent trials) or push (alcohol-incongruent trials) (*F*_(1,74)_ = 22.81, *p* < 0.001), with more errors observed during push responses than pull responses (mean difference = 0.205, t(74) = 4.03, p < 0.001). Group × Action interaction was found to be non-significant (F(1,74) = 0.378, p = 0.541). Together, these results suggest that alcohol cues generally biased participants toward approach-consistent responses, leading to greater difficulty when participants were required to execute avoidance (push) actions, regardless of their automatic approach or avoidance tendencies toward alcohol.

Although the Group × Action interaction was not significant at the behavioral level, ERP analysis revealed automatic tendency-based differences in neural processing during the PIT task. A significant interaction was observed for the N2 component at right-frontal sites (*F*_(1,37)_ = 69.637, *p* < 0.001; Figure 3B), whereas no such effect was evident for the centroparietal P3 (*F*_(1,37)_ = 0.125, *p* = 0.726, *BF_10_* ≈ 0.106; Figure 3C). The N2 is commonly associated with cognitive control processes such as response selection and the regulation of competing action tendencies. Interestingly, the approach group showed significantly reduced N2 amplitudes during push actions compared to the avoidance group (median difference = 2.500, *W* = 274, *p* < 0.002, r = 0.600; Figure 3B), suggesting diminished engagement of frontal control mechanisms when Pavlovian alcohol cues conflict with avoidance responses.

**Figure 3.**
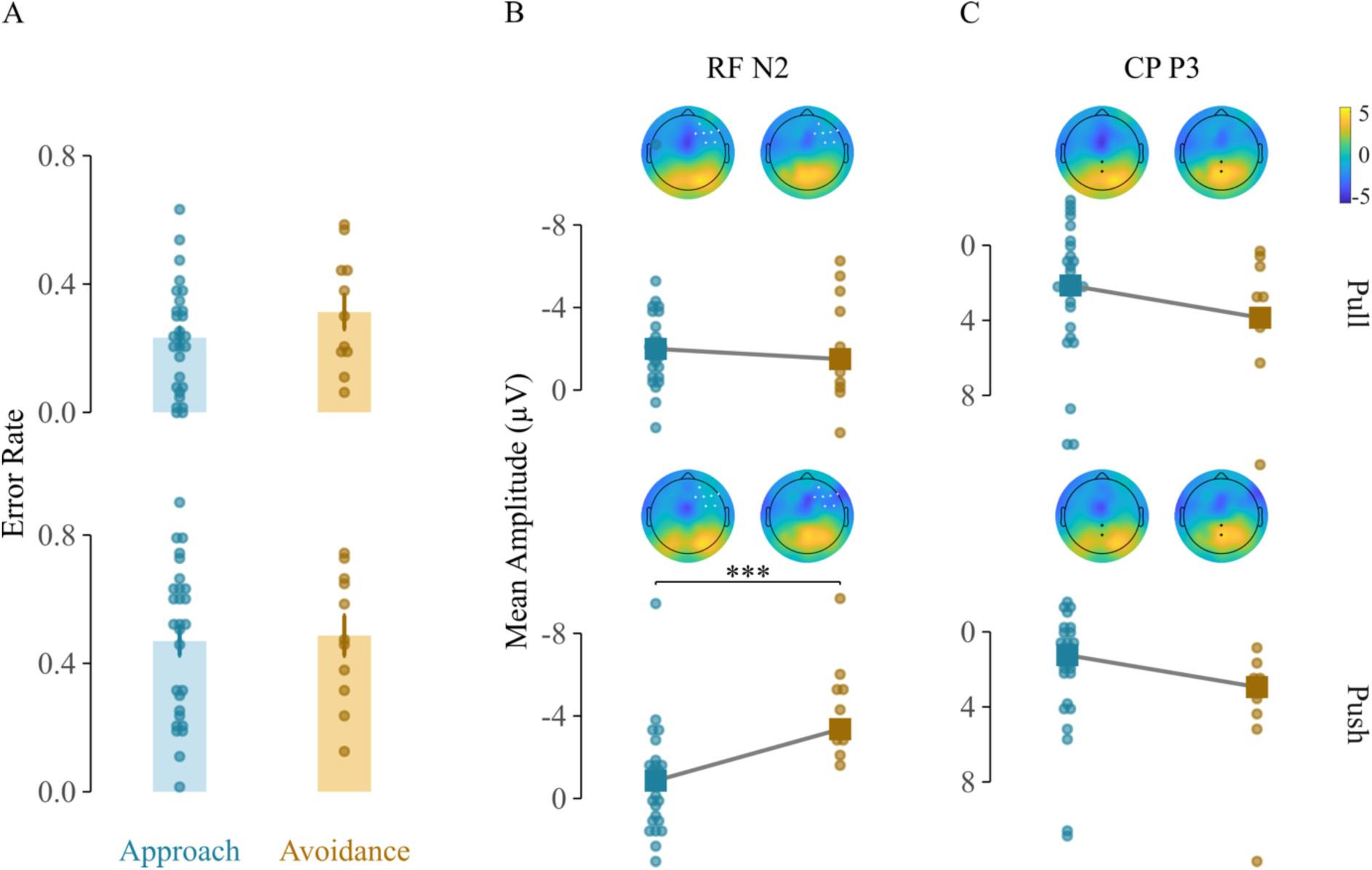
The approach group shows lower interference from alcohol cues during pull responses after conditioning (during baseline PIT, Panel A), with a corresponding reduction in neural response at the RF during push responses (Panel B). However, no such change was evident at CP (Panel C). Blue and brown colors indicate approach and avoidance group datapoints, respectively. RF = right frontal region; CP = centro-parietal region. * p < 0.05; ** p < 0.01; *** p < 0.001

#### Counterconditioning-Based Modulations Across Groups

Counterconditioning (CC) targets maladaptive stimulus–outcome associations by pairing conditioned stimuli with outcomes of opposing valence, thereby updating appetitive associations. In the current study, participants viewed alcohol cues paired with loss outcomes across multiple trials, aiming to reverse previously acquired alcohol-reward expectancies toward negative outcome representations. Following CC of the alcohol cues, both the approach and avoidance groups associated alcohol cues with negative outcomes, with no significant group differences (*χ²*_(1, N=39)_ = 0.01, *p* = 0.92), confirming that CC similarly altered participants’ explicit associations across groups.

#### Counterconditioning Selectively Reduced Automatic Approach Bias and Enhanced Neural Control During A-AAT

Despite equivalent explicit learning, CC produced differential effects on AAI and respective neural correlates depending on participants’ pre-existing automatic tendencies. A significant Group (Approach vs. Avoidance) × Assessment (Baseline vs. No R-Cue Post CC vs. R-Cue Post CC) interaction (F(2,74) = 11.843, p < 0.001; Figure 4A) indicated that CC differentially influenced AAI across groups. AAI was selectively reduced in the approach group, both during trials with (median difference = −0.098, W = 281, p < 0.001, r = 0.584) and without retrieval cues (median difference = −0.085, W = 324, p = 0.002, r = 0.404), where presence of retrieval cues produce an additional reduction over no-cue trials (median difference = −0.013, W = 281, p = 0.028, r = 0.212). The avoidance group showed no significant AAI change under either condition (BF₁₀ ≈ 0.835 and 0.611, respectively). Notably, even after CC, AAI in the approach group remained elevated above the avoidance group baseline levels (median difference = −0.068, W = 238, p = 0.009, r = 0.420), indicating a need for further reduction in automatic approach tendencies.

**Figure 4.**
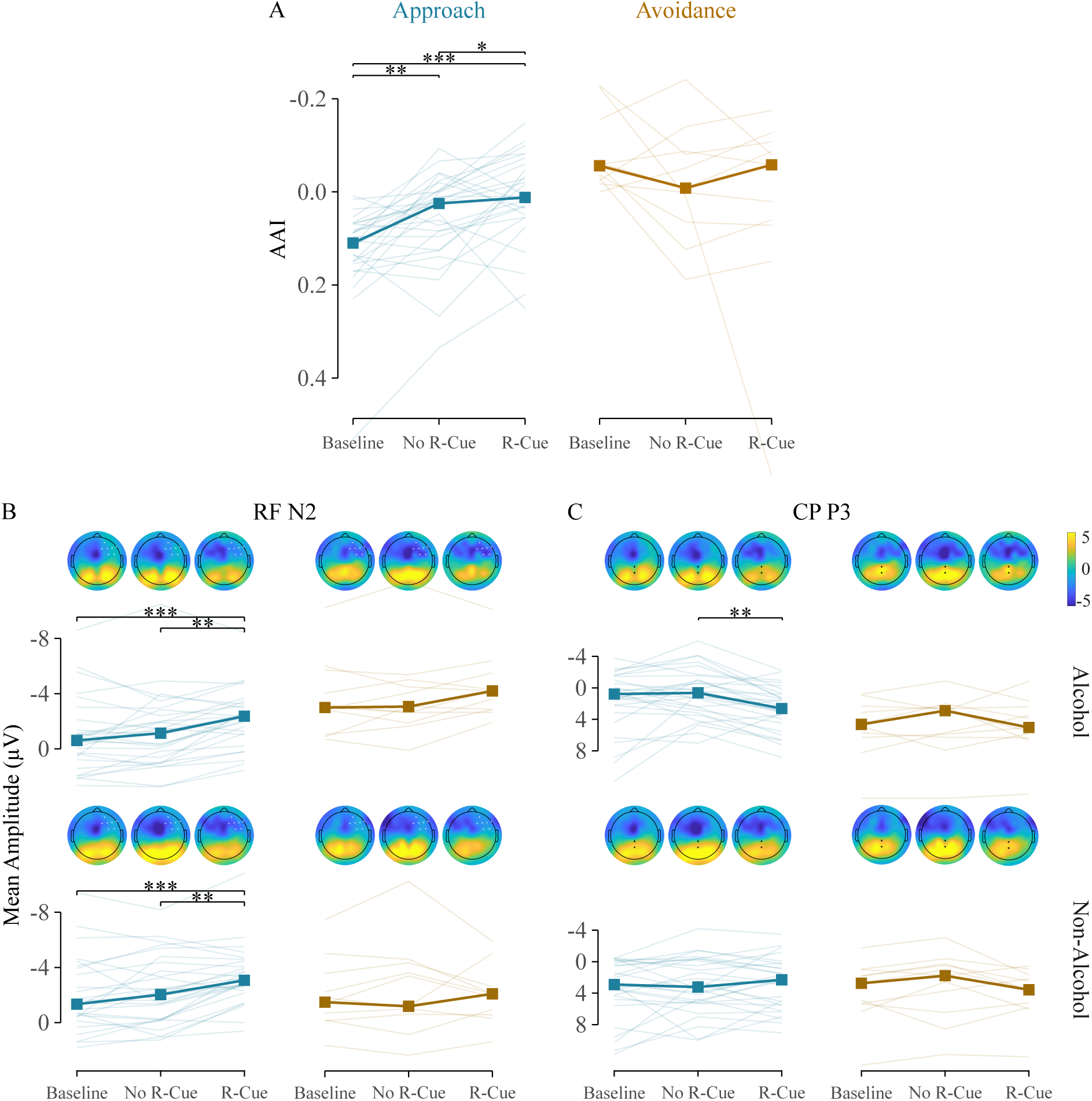
After CC, individuals with approach tendencies showed significant reductions in automatic approach tendency towards alcohol cues (Panel A), along with neural enhancements at both RF (Panel B) and CP (Panel C), particularly in the presence of a retrieval cue. Baseline: baseline PIT response (before CC); No R-Cue = no retrieval cue trails post-CC; R-Cue = retrieval cue trails post-CC; RF = right frontal region; CP = centro-parietal region. * p < 0.05; ** p < 0.01; *** p < 0.001

##### Right Frontal N2

Changes in AAI after CC were accompanied by enhanced N2 amplitudes at RF (Group x Assessment: F_(2,407)_ = 43.153, p < 0.001; Figure 4B). Particularly, CC enhanced N2 amplitudes in the approach group when retrieval cues were present, for both alcohol (median difference = 1.756, W = 39, p < 0.001, r = 0.706) and non-alcohol stimuli (median difference = −1.729, W = 40, p < 0.001, r = 0.701), with no enhancement in the absence of retrieval cues (BF₁₀ ≈ 0.343 and 0.445, respectively). The retrieval cue effect, compared to no retrieval cue trials, was significant across both stimulus types (alcohol: median difference = −1.230, W = 61, p = .001, r = .611; non-alcohol: median difference = −1.036, W = 57, p = 0.001, r = 0.629; Figure 4C), indicating that retrieval cues enhanced early frontal neural responses regardless of stimulus category. The avoidance group showed no N2 changes following CC (BF₁₀ ≈ 0.341 and 0.491).

##### Centroparietal P3

A significant Group × Assessment interaction was also observed for P3 amplitudes (F(2,407) = 14.840, p < 0.001; Figure 4C). In the approach group, P3 remained largely unchanged from baseline in both post-CC conditions (BF₁₀ ≈ 0.805 and 0.561), but was significantly enhanced on alcohol trials when retrieval cues were present versus absent (median difference = −1.986, W = 358, p = 0.001, r = 0.667; Figure 4C), indicating retrieval-cue-dependent engagement of attentional processing. The avoidance group showed no P3 modulation under either condition (BF₁₀ ≈ 0.356 and 0.311). This indicates that retrieval cues were associated with greater attentional allocation to alcohol stimuli in the approach group, but not in the avoidance group.

### Counterconditioning Strengthened Avoidance-Congruent Instrumental Responding During PIT

Beyond CC’s effects on automatic bias and neural responses during the A-AAT, it also reshaped how alcohol cues influenced instrumental action execution during the PIT task. CC significantly modulated both error rates during the PIT task (Group × Action × Assessment: F(3,259) = 19.481, p < .001; Figure 5A), indicating that the effects of CC on cue-driven action tendencies were behaviorally expressed. In the approach group, alcohol cue–induced interference during push trials decreased significantly from baseline only when retrieval cues were present (median difference = 0.438, W = 336, p = 0.004, r = 0.570), not without them (BF₁₀ ≈ 0.276), consistent with CC degrading approach-biasing expectancies. Conversely, the presence of retrieval cues increased pull interference in the approach group (median difference = −0.408, W = 62, p = .004, r = −.574; Figure 5A). Taken together, this opposing pattern across action types, where CC with retrieval cues reduced push interference but increased pull interference, suggests that CC strengthened avoidance responses while making approach responses more difficult to execute. In the avoidance group, no behavioral changes were observed after CC (all p > 0.05).

**Figure 5.**
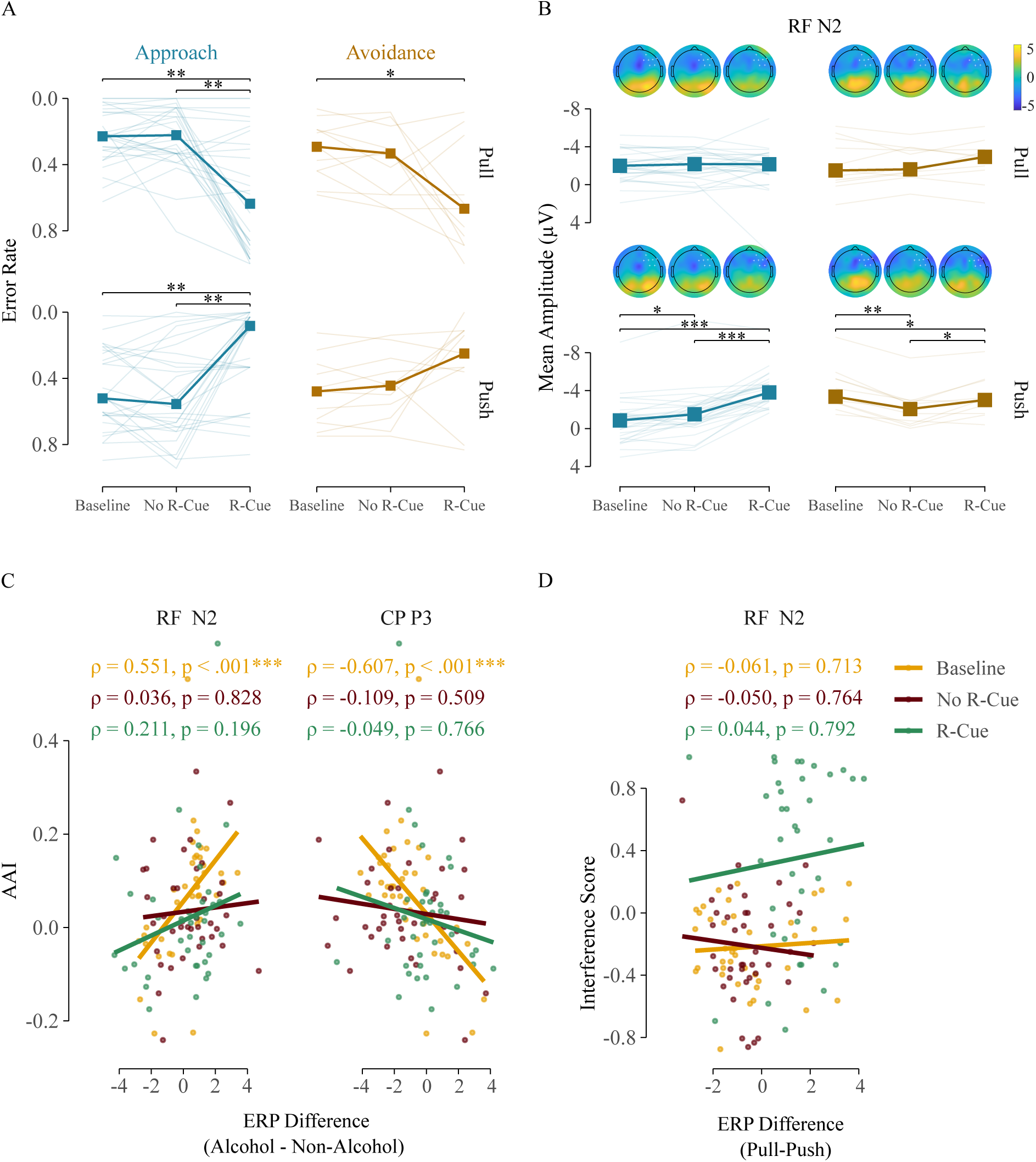
The approach group shows prominent changes after CC, with enhanced push actions (lower error rates) in the presence of a retrieval cue, whereas both groups exhibit increased interference during pull actions (Panel A). Push actions were further complemented by enhanced neural response at RF in the approach group (Panel B). During the baseline, a significant relationship was observed between AAI and ERP difference scores, which later disappeared after CC (Panel C). However, no significant association emerged between behavioral and neural responses during PIT (Panel D). Baseline: baseline PIT response (before CC); No R-Cue = no retrieval cue trails post-CC; R-Cue = retrieval cue trials post-CC; AAI = approach avoidance index; RF = right frontal region; CP = centro-parietal region. * p < 0.05; ** p < 0.01; *** p < 0.001

CC also significantly modulated RF N2 amplitudes during the PIT task (F(3,259) = 10.247, p < 0.001; Figure 5B). In the approach group, the behavioral improvement in push trials was accompanied by enhanced RF N2, with retrieval cue trials showing a larger effect (median difference = 2.950, W = 403, p < 0.001, r = 0.712) than non-retrieval trials (median difference = 0.640, W = 317, p = 0.016, r = 0.191). This retrieval cue–dependent N2 enhancement was confirmed by direct comparison between no-retrieval and retrieval cue trials post-CC (median difference = 2.310, W = 9, p < 0.001, r = 0.646; Figure 5B). No corresponding neural changes were observed for pull trials in the approach group (all p > 0.05). In the avoidance group, despite unchanged behavior, RF N2 was enhanced during push responses on both non-retrieval (median difference = −0.342, W = 7, p = 0.038, r = 0.203) and retrieval trials (median difference = −1.299, W = 0, p = 0.006, r = 0.483), with retrieval cues producing a significantly larger enhancement compared to no-retrieval cue trials (median difference = 0.957, W = 5, p = 0.030, r = 0.357). Together, these results indicate that CC was associated with enhanced RF N2 during push responses, particularly in retrieval cue trials, with corresponding behavioral improvement in the approach group but not in the avoidance group.

### Counterconditioning Dissociated Neural–Behavioral Coupling During A-AAT

To assess the specificity of CC’s effects, we examined whether the baseline relationship between automatic tendencies and neural responses was altered post-CC. At baseline, AAI was significantly correlated with ERP amplitude differences between alcohol and non-alcohol cues (N2: ρ = 0.551, p < .001; P3: ρ = −0.607, p < 0.001; Figure 5C), indicating that stronger approach biases were associated with attenuated neural responses to alcohol stimuli. Following CC, these correlations were no longer significant under either condition (all p > 0.05; Figure 5C). To confirm a true change, correlation coefficients were directly compared using Steiger’s tests. For RF N2, no significant differences emerged across any assessment phase (all p > 0.05), indicating that CC did not reliably alter the relationship between approach tendencies and early frontal processing. For CP P3, the baseline AAI–P3 correlation differed significantly from the retrieval cue condition (t(36) = −2.67, p = 0.01), as did the no retrieval cue versus retrieval cue comparison (t(36) = −2.58, p = 0.01), whereas baseline and no retrieval cue conditions did not differ (t(36) = −0.16, p = 0.87). This pattern demonstrates that CC selectively abolished the coupling between approach tendencies and P3-indexed attentional processing specifically in the presence of retrieval cues.

For PIT, no significant correlations were observed between N2 difference scores and behavioral interference at baseline or after CC (all p > 0.05; Figure 5D), suggesting that neural and behavioral measures during PIT capture partially dissociable aspects of cue-driven interference.

To examine whether post-CC changes in PIT interference were independently predicted by concurrent changes in approach tendencies, PIT change scores were regressed on AAI change scores separately for retrieval cue and no-cue conditions. Neither pull (retrieval cue: b = 0.105, p = 0.849; no cue: b = −0.130, p = 0.625) nor push (retrieval cue: b = −0.178, p = 0.669; no cue: b = 0.155, p = 0.589) interference changes were significantly predicted by AAI change under either condition, a result that remained consistent when robust HC3 standard errors were applied. Spearman correlations further confirmed the absence of reliable associations (all p > 0.05). These findings indicate that CC-induced modulation of instrumental responding occurred independently of concurrent reductions in approach bias, suggesting that PIT changes reflect the direct effect of CC on cue-outcome expectancies rather than a byproduct of approach bias reduction.

## Discussion

The present study examined how automatic approach and avoidance tendencies toward alcohol modulate Pavlovian-to-instrumental transfer (PIT), and whether counterconditioning (CC) combined with retrieval cues can attenuate automatic approach biases and cue-driven instrumental interference. Approach-oriented individuals showed attenuated right frontal N2 during push actions despite comparable behavioral PIT performance across groups, indicating that automatic tendencies compromise early frontal processing during incongruent instrumental responding without necessarily manifesting as overt behavioral differences. CC reduced automatic approach biases and recalibrated cue-driven instrumental interference in the approach group, reflected in reduced push interference and increased pull interference, accompanied by restoration of previously attenuated neural activity. Importantly, CC-induced changes in the alcohol approach index did not predict PIT interference, suggesting that CC affected these two processes through partially dissociable pathways.

### Individual Differences in Automatic Tendencies Shape Neural Processing During Instrumental Behavior

While prior PIT investigations have predominantly examined alcohol cue interference without considering how opposing automatic tendencies might produce distinct interference patterns (Hogarth et al., 2019; Seabrooke et al., 2019; Watson et al., 2014), our findings reveal that approach and avoidance tendencies produced qualitatively different neural signatures during instrumental responding. Behaviorally, error rates were comparable across groups, but approach-oriented individuals showed attenuated right frontal N2 during push actions, suggesting reduced engagement of early frontal regulatory processes when instrumental avoidance actions conflicted with prepotent stimulus-driven tendencies (Botvinick et al., 2001; Sharbanee et al., 2014). This behavioral-neural dissociation suggests that strong learned associations between alcohol cues and rewarding outcomes reduce frontal recruitment in response to alcohol stimuli (Stacy & Wiers, 2010; Verma et al., 2025; Wiers et al., 2008), leaving individuals neurally under-equipped to manage cue-action incongruence even when overt performance appears intact. The absence of centroparietal P3 differences suggests this compromise was specific to early frontal processing, consistent with later-stage attentional processing being less sensitive to Pavlovian interference when tasks do not require explicit cue evaluation (Nadler et al., 2011).

The comparable behavioral interference across both actions in the avoidance group highlights a boundary condition of dual-process models, which capture the facilitatory influence of approach biases but do not readily account for avoidance-oriented behavior. Avoidance likely reflects the relative absence of a strongly rooted approach drive rather than an opposing bias of equivalent strength (Carver, 2006), suggesting that the neural signature of PIT depends on the strength of underlying AAT profiles. The attenuation of N2 in approach-oriented individuals is consistent with strong cue-driven tendencies reducing frontal regulatory engagement, i.e., when an alcohol cue strongly primes approach, push responses fail not through heightened neural effort but through insufficient recruitment of the frontal processes supporting response regulation (Buss et al., 2011; Espinet et al., 2012). These findings demonstrate that individual differences in automatic response tendencies shape the neural underpinnings of cue-driven interference even when behavioral outcomes appear comparable, adding complexity to existing accounts of habitual and goal-directed processes in PIT.

### Counterconditioning Attenuates Approach Tendencies and Recalibrates Cue-Driven Interference Through Partially Dissociable Mechanisms

CC established competing associations that altered what alcohol cues signal, weakening their capacity to bias instrumental behavior toward approach-congruent actions (Hogarth et al., 2014; Houben et al., 2012; Wiers et al., 2011), consistent with CC operating by directly modifying the associative basis of approach biases rather than relying on effortful top-down control (Baker & Cannon, 1979; Das et al., 2015; Goltseker et al., 2016, 2021). It is worth considering why CC can be effective in the laboratory when naturally occurring negative consequences of alcohol use, such as health problems, relationship damage, and financial loss, often fail to produce similar changes in real-world drinking behavior. Primarily, immediate positive reinforcement from alcohol occurs within minutes, whereas negative consequences are typically delayed by hours or days, making temporal contiguity poor in naturalistic settings. Real-world negative consequences are also probabilistic and inconsistent, and are frequently attributed to factors other than alcohol itself, preventing direct associative learning. Laboratory CC addresses these issues by providing immediate, consistent, and unambiguous cue-aversive outcome pairings that naturalistic experience cannot achieve.

Critically, CC-induced reductions in approach bias did not predict changes in PIT interference. This independence suggests that CC simultaneously but separately modified two distinct processes: the appetitive cue-outcome associations underlying approach bias, and the expectancy structure sustaining cue-driven instrumental interference. This is broadly consistent with Hogarth et al.’s (2014) demonstration that PIT effects are specifically sensitive to the degradation of instrumental expectancies and that procedures targeting cue-outcome associations alone are insufficient to abolish cue-driven instrumental responding. CC may therefore have reduced approach bias by reassigning the aversive value of alcohol cues, while its effect on PIT operated through a related but distinct pathway involving degradation of cue-based action selection signals.

The specificity of CC effects to the approach group is consistent with its mechanism of action. CC targets existing appetitive associations, and individuals with stronger baseline approach biases provided a richer substrate for modification. Avoidance-oriented individuals, lacking strong appetitive alcohol associations, showed minimal change, consistent with CC modifying existing learned associations rather than establishing entirely new response orientations. This pattern suggests that individuals with stronger approach biases are both more susceptible to cue-driven neural disruption during instrumental responding and more responsive to CC.

### The Role of Retrieval Cues in Facilitating CC Outcomes

Retrieval cues enhanced CC outcomes by enabling reactivation of newly formed aversive associations at test. Consistent with Hogarth et al.’s (2014) demonstration that discriminative procedures degrading cue-outcome expectancies were more effective than simple extinction in abolishing PIT, retrieval cues may have provided a contextual scaffold supporting expression of competing associations across contexts (Collins & Brandon, 2002; Stasiewicz et al., 2007; Treanor et al., 2017), consistent with evidence that interventions incorporating retrieval cues produce stronger and more persistent effects (Lörsch et al., 2024).

However, alternative mechanisms warrant consideration. Retrieval cues were paired with larger monetary losses during CC and may therefore have formed direct aversive associations independently of their link to alcohol cues, supported by the non-alcohol-specific enhancement of frontal N2 when retrieval cues were present. Additionally, the consistent co-presentation of retrieval cues and alcohol cues may have produced compound stimulus encoding rather than purely contextual reminding. Future designs should present retrieval cues in isolation during test phases to disentangle whether they reactivate specific alcohol-loss associations, function as direct aversive cues, or create compound representations. Despite these ambiguities, retrieval cues consistently enhanced CC outcomes behaviorally and neurally.

### Neural Mechanisms Underlying Approach Tendency Modulation and CC-Induced Recalibration

At baseline during the A-AAT, approach-oriented individuals showed attenuated N2 and P3 amplitudes at right frontal and centroparietal regions in response to alcohol relative to non-alcohol cues, reflecting reduced engagement of early evaluative and attentional processes triggered by alcohol stimuli (Buss et al., 2011; Espinet et al., 2012; Polich, 2007; Reed et al., 2022), consistent with more automatic stimulus-driven processing. The convergence of attenuated N2 and P3 in the A-AAT, compared to only N2 attenuation in the PIT, suggests that task demands modulate the degree of neural compromise. The A-AAT places concurrent demands on both early and late-stage regulatory processes through explicit cue-based responding, whereas PIT selectively engages early frontal processing, given that cues influence behavior implicitly rather than through explicit evaluation.

Following CC, previously attenuated N2 and P3 amplitudes were restored in the approach group. Enhanced N2 during push actions in the PIT is consistent with CC weakening automatic cue-approach associations that would otherwise suppress frontal regulatory recruitment during incongruent responding (Das et al., 2015; Van Gucht, Baeyens, et al., 2010). The disruption of the AAI-ERP correlation following CC, particularly for P3 under retrieval cue conditions, suggests that attentional responses to alcohol cues became decoupled from approach bias strength following the establishment of competing aversive associations. That P3 modulation was less consistent in the PIT context is consistent with CC primarily targeting automatic associative processes rather than deliberate attentional control(Das et al., 2015; Kerkhof et al., 2011). The independence of neural and behavioral PIT measures across all phases further indicates that right frontal N2 and behavioral error rates capture partially dissociable aspects of cue-driven interference.

### Practice Effects as an Alternative Explanation

Post-CC changes could alternatively reflect practice or repeated-measures effects. However, several observations argue against this. First, effects were largely specific to the approach group, whereas practice effects would be expected across both groups. Second, retrieval cue conditions showed systematically larger changes than no-cue conditions, inconsistent with generic task repetition. Third, the direction of changes was also theory-consistent, with reduced approach bias, restored neural responses, and opposing modulation of pull versus push interference. Nevertheless, without a no-CC control group receiving repeated task administrations without CC, practice contributions cannot be definitively excluded. Future studies should include such conditions to isolate CC-specific effects.

### Limitations and Future Directions

Several limitations warrant acknowledgment. First, all tasks were completed within a single session, limiting findings to immediate pre-consolidation processes. Results should therefore be interpreted as proof-of-concept for short-term associative modification, with future multi-session research needed to examine consolidation and persistence of effects. The single-session design also precludes examination of reconsolidation-based memory updating (Soeter & Kindt, 2015), which may be necessary for more durable change. Second, the unequal group sizes (approach n = 28; avoidance n = 11), arising from post-hoc classification, reduced statistical power for avoidance group comparisons, as reflected in inconclusive Bayes factors and borderline p-values. Preliminary screening in future studies should ensure more balanced representation. Third, the use of identical alcohol stimuli across baseline, conditioning, and CC phases also means that stimuli acquired multiple potentially conflicting associations, and observed PIT changes may partly reflect degradation of original conditioning rather than purely competing association formation. Multi-session designs separating these phases would help disentangle these contributions. Finally, the subclinical sample limits generalizability to alcohol use disorder, where more entrenched associative learning, altered neurobiology, and clinical craving may produce qualitatively different effects. Replication in clinical populations is essential for determining the therapeutic utility of CC.

### Conclusion

This study establishes that automatic approach tendencies toward alcohol shape early frontal neural processing during instrumental responding, with approach-oriented individuals showing attenuated right frontal activity during incongruent push actions despite comparable behavioral performance. CC combined with retrieval cues reduced automatic approach biases and recalibrated cue-driven instrumental interference, restoring previously attenuated neural activity in approach-oriented individuals. Importantly, CC-induced changes in approach bias did not predict PIT interference, indicating that CC affected these processes through partially dissociable pathways rather than a single unified mechanism. These findings advance understanding of how individual differences in automatic response tendencies shape Pavlovian-instrumental interactions, and underscore the importance of accounting for individual tendency profiles when designing and evaluating associative interventions for alcohol-related behavior. Future research in clinical populations across multiple sessions is needed to determine whether these mechanisms contribute to pathological alcohol use and whether CC can produce clinically meaningful and lasting change.

## DECLARATIONS

### Conflict of Interest

None

### Data Availability

The datasets used and/or analyzed during the current study are available from the corresponding author upon reasonable request.

## Acknowledgement

This work was supported by the Ministry of Education, Government of India, through the Prime Minister’s Research Fellowship (PMRF) awarded to AKV.

## Notes

### Competing Interest Statement

The authors have declared no competing interest.

### Summary of Updates

This version clarifies the background, reorganizes the results into a more coherent logical flow, and revises the discussion accordingly for improved clarity and interpretation.

